# A molecular approach to studying Hymenoptera diets using polistine wasps

**DOI:** 10.1101/2020.04.06.024422

**Authors:** M.-C. Lefort, J.R. Beggs, T.R. Glare, E.J. Doyle, T.E. Saunders, S. Boyer

## Abstract

- The study of animal diets has benefited from the rise of high-throughput DNA sequencing applied to stomach content or faecal samples. The latter can be fresh samples used to describe recent meals, or older samples, which can inform about past feeding activities. For most invertebrates, however, it is difficult to access ‘historical’ samples, due to the small size of the animals and the absence of permanent defecation sites. Therefore, sampling must be repeated to account for seasonal variation and to capture the overall diet of a species.
- This study develops a method to describe the overall diet of nest-building Hymenoptera based on a single sampling event, by analysing prey DNA from faeces accumulated in brood cells. We collected 48 nests from two species of introduced paper wasps (*Polistes chinensis*, and *P. humilis*) in the urban and peri-urban areas of Auckland, New Zealand, and selected two samples per nest. One from brood cells in the outer layer of the nest to represent the most recent diet, and one from brood cells in an inner layer to represent older diet.
- Diet differed between species, although both fed mainly on Thysanoptera, Lepidoptera and Acariformes. Prey taxa identified to species level included both agricultural pests and native species. Prey communities consumed were significantly different between inner and outer nest samples suggesting seasonal variation in prey availability and/or a diversification of the wasps’ diet as the colony grows. We also show for the first time potential predation of marine organisms by Polistes wasps.
- Our study provides field evidence that polistine wasps feed on agricultural pests, supporting the hypothesis that some social wasp species could have a suppressing effect on agricultural pests. The proposed methodology is readily applicable to other nest-building Hymenoptera and has the potential to provide comprehensive knowledge about their diet with minimum sampling effort. Such knowledge is Essential to measure the ecological impact of invasive Vespidae and support the conservation of native invertebrate biodiversity.

## 1 INTRODUCTION

In recent years, the study of invertebrate diets has been improved through the application of molecular methods to detect trophic interactions (Sheppard et al. 2005; Boyer et al. 2013; González-Chang et al. 2016). These methods allow the sequencing of prey DNA present in the regurgitates (Waldner & Traugott 2012), gut contents (Olmos-Pérez et al. 2017) or faeces (Boyer et al. 2011) of predators. Because these methods analyse fresh material taken from live individuals, the samples essentially provide a snapshot of recently consumed prey items based on DNA still present in the gut at the time of capture. Understanding temporal variation in gut contents is essential to characterise an animal’s diet, but such data is difficult to collect, and therefore under-represented in food web studies (McMeans et al. 2019). Nevertheless, many taxa exhibit significant seasonal variation in their diets (Waterhouse et al. 2014; Lambert & Rothman 2015; Coulter et al. 2019; Amponsah-Mensah et al. 2019). Recent developments in molecular techniques mean that even low-quantity and low-quality DNA samples can be analysed efficiently, and applying these techniques to the analysis of faecal samples for assessing diet represents an interesting opportunity (Monterroso et al. 2019, Waterhouse et al. 2014). This could be a particularly useful strategy for assessing seasonal variation in the diets of taxa who deposit their faeces in permanent sites such as latrines, or other territory marking sites (Fretueg et al. 2015). Such approach could provide valuable insight to the diet of nest-building Hymenoptera where larval faeces accumulate within the nest.

Social wasp colonies are organised around a caste system containing a reproductive queen, male drones, and female workers which construct a nest out of mud or wood fibres (Oster & Wilson 1978). During their development, the larvae of social wasps are fed a range of food by the workers, including other invertebrates, by adult wasps (Harris & Oliver 1993; Kasper et al. 2004; Todd et al. 2015). Larvae remain in individual cells in the nest until metamorphosis, while their faeces (or frass) accumulate at the bottom of their brood cell. This frass constitutes a historical record of the animal’s diet throughout its development and, in multi-layered nests, we hypothesize that faeces in successive layers of brooding cells could be used to identify which prey species adult wasps have brought to their young over several weeks or months. Social wasps are considered pests in many regions of the world (Beggs et al. 2011; Lester & Beggs 2019), particularly species of Vespula and Polistes introduced into Australasia, Hawaii and South America (Clapperton et al. 1996; Matthews et al. 2000; Masciocchi & Corley 2013; Hanna et al. 2014). For example, in New Zealand, introduced Vespula cause significant impacts on the biodiversity and ecology of beech forest ecosystems (Moller et al. 1991; Beggs et al. 2005) by monopolising the honeydew produced by native scale insects (Hemiptera: Sternorrhyncha) (Beggs & Wardle 2006). Through their feeding on this abundant resource, introduced wasps are able to attain their highest densities in the world and, in doing so, compete for food resources with many native species including birds, bats, lizards and other native insects that are known to feed on honeydew (e.g. Harris 1991; Toft & Rees 1998; Beggs 2001; Elliott et al. 2010). Each year, social wasps cause millions of dollars of damage to the New Zealand economy, primarily due to the impact of wasps on honeybees (*Apis mellifera* L.) and the loss of flow-on benefits to pastoral farming (MacIntyre & Hellstrom 2015b). Beyond their economic and ecological impacts, wasps cause considerable nuisance problems through their painful and occasionally life-threatening stings (Golden et al. 2006). While the risks posed by these species are widely known, their ecological roles may be more complex than is currently recognised, particularly if they replace functional diversity lost through human impacts (Beggs & Wardle 2006). Additionally, social wasps, and in particular *P. chinensis*, may contribute biocontrol services in agricultural systems as generalist predators of the main insect orders of crop pests, Lepidoptera and Diptera (Todd et al. 2015; Southon et al. 2019). A more comprehensive analysis of the diets of social wasps would allow a more nuanced understanding of the ecological roles these insects play.

The main aim of the study was to develop a new molecular method to study the diet of social wasps by retrieving prey DNA from faeces left by wasp larvae inside the nest. Historically, wasp diets have been studied by collecting and identifying the prey items carried by adult foraging wasps when they return to the nest to feed the larvae (Kasper et al. 2004) or by dissecting the guts of adult wasps (e.g. Ward & Ramón-Laca 2013). These traditional methods only provide a snapshot of the diet of the insect at a given point in time, while the method we propose here offers an overview of the diet of the colony throughout the lifespan of the nest. We apply our method to two sympatric polistine wasps to examine the way they partition resources in urban and peri-urban habitats in New Zealand, and to better understand their ecological roles in relation to New Zealand invertebrates.

## 2 MATERIALS AND METHODS

### 2.1 Nest monitoring and sampling

We collected polistine nests in the Auckland region, New Zealand, during the peak of the 2017 summer season (i.e. January). Potential nesting sites were located with the aid of previous occurrence records on iNaturalistNZ (URL: https://inaturalist.nz/), an open biodiversity observation platform built to record the occurrence of taxa across New Zealand. A total of 53 active nests were collected between 1st of March and 15th of May 2017 (Figure 1). After removing resident adults from the nest with commercial fly spray, nests were placed in individual sealable plastic bags and into a cold storage container for transportation, before being stored at −80°C for optimal DNA preservation.

**FIGURE 1.**
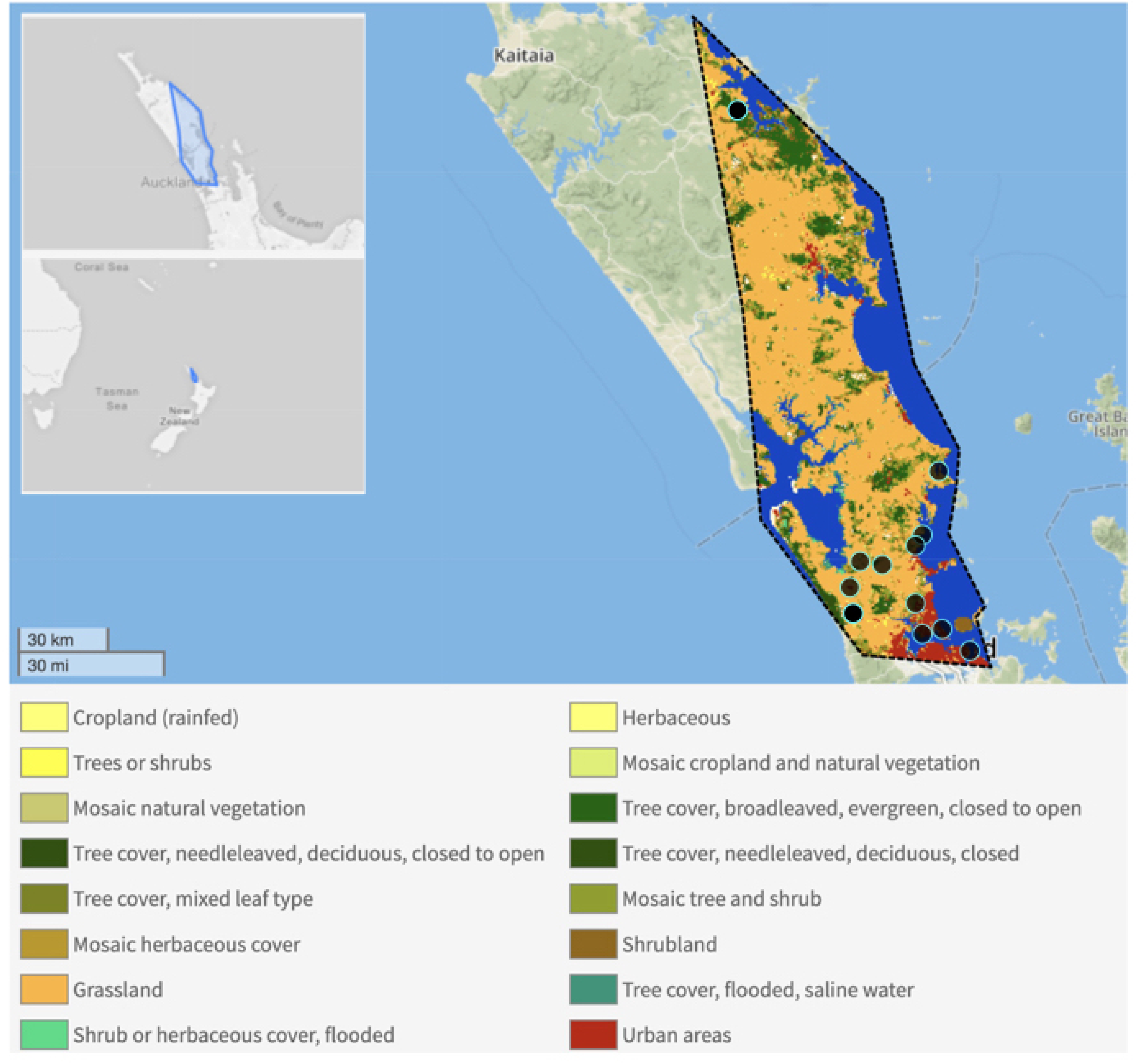
Location of samples collected in and around Auckland. Black dots correspond to areas where one or several nests were collected. See legend for colour code for land cover classification (from geofolio.org).

### 2.2 DNA extractions

Among the collected nests, only the largest and best preserved were used for analyses (n=44 for *P. chinensis* and n=4 for *P. humilis*). Using bleached and sterilised tweezers, insect frass was sampled from one cell located on the outer (top) layer and from one cell located on the inner (bottom) layer of each nest. These two layers represent different batches of brood, and therefore they preserve a record of the diet of the colony at two different times. Faecal samples were ground directly in microcentrifuge tubes using a hand-held mortar and pestle and the DNA was extracted using a ZR Tissue & Insect DNA MicroPrep extraction kit (Zymo, Irvine CA) following an existing methodology for low quantity and degraded insect environmental DNA (Lefort et al. 2012). DNA concentration in the resulting eluates was measured by fluorometry using a Qubit®dsDNA HS Assay Kit (Life Technologies, Carlsbad, CA).

### 2.3 DNA amplification and sequencing

Because prey DNA was likely to be highly degraded, primers targeting a short (313 bp) gene region of the mitochondrial gene COI were used for amplification. We selected the pair of primers mICOIintF (forward GGWACWGGWT-GAACWGTWTAYCCYCC) and HCO2198 (reverse TAAACTTCAGGGTGACCAAAAAATCA), which are recommended for broad coverage of metazoan biodiversity, (Leray et al. 2013; Lear et al. 2018). PCR reactions comprised 1 mL BSA, 10 mL Green GoTaq Mix, 0.6 μL of each primer [10 μM], 5.8 μL of ultrapure water, and 2 μL of template. A few recalcitrant samples were amplified using less template (0.5-1 μL) to limit the effect of potential PCR inhibitors. We used the touchdown PCR protocol recommended by Leray et al. (2013), with a 2 min initial denaturation (95°C) step. After the initial denaturation we conducted 16 cycles of 10 s denaturation (95°C), 30 s annealing (62°C -1°C per cycle) and 60 s elongation (72°C), followed by 25 cycles where the annealing temperature remained constant (46°C), and a final 7 min elongation step was then carried out (72°C). According to standard protocol (Support Illumina 2016), all PCR products were purified using AMPure magnetic beads (Agencourt) and DNA concentration was standardised to 2 μM in all samples before pooling for high-throughput DNA sequencing. The resulting library was processed in high-throughput DNA sequencing analysis on one run of Illumina MiSeq using the 300 × 300 paired end protocol as recommended by the manufacturer. Ligation of individual barcodes, pooling of the libraries, and sequencing, were performed by New Zealand Genomics Ltd., Auckland, New Zealand.

### 2.4 Data analysis

The bioinformatic workflow was perfomed by NGBS (Nextgen Bioinformatic Services, New Zealand), based on the *vsearch* pipeline (Rognes et al. 2016) and included quality control, merging of paired end reads, removal of singletons, dereplication, and chimera detection using *uchime*. The merged reads were then quality filtered and clustered at 97% identity to generate Molecular Operational Taxonomic Units (MOTUs). Following de novo and reference-based chimera detection, a final number of 1,436 MOTUs were detected, with an average read length of 365 bp (close to the expected amplicon size). Each MOTU was then compared to the NCBI NT database using BLASTn. To limit false positives and remove potential sequencing errors, only MOTUs for which read abundance within a sample was superior to 0.5% of the total were retained. To ensure reliable identification, only sequences for which the best hit had a query coverage over 70% were retained, which corresponded to an overlap of 250 bp or more between the query and the best hit. Retained MOTUs were identified to species level when the percentage identity was ≥ 98%, or assigned to the order of the corresponding best hits when their percentage identity was between 80 and 98%. Reads with percentage identity < 80% were not retained for prey identification. MOTUs for which the best hit had more than 98% identity, but the species name was not available in Genbank (e.g. only genus or family name), were searched against the Barcode Of Life Database (BOLD). Only MOTUs identified as terrestrial invertebrates were considered as prey. Reads corresponding to *Polistes* wasp DNA were used to confirm species identification of the wasp to which the nest belonged. Prey MOTUs confidently identified at the species level were categorised as native or introduced species based on Scott (1984) and the New Zealand Organisms Register (http://www.nzor.org.nz/).

### 2.5 Statistical analysis

Because of an imbalance in the number of samples analysed for (n = 8) and *P. chinensis* (n = 88), we used the non-parametric Kruskall-Wallis test to compare the number of reads as well as prey richness between the two wasps species. To compare between inner and outer samples, we used the non-parametric Wilcoxon test to account for the non-normality and paired nature of the data. The cumulative number of MOTUs detected from the inner and outer layer of *P. chinensis*’ nests was compared using a Koglomorov-Smirnov test. Because the analysis of only eight samples from four nests for *P. humilis* led to a low coverage of its diet, results for this species are only considered as indicative and a detailed analysis of prey community was only conducted for *P. chinensis*. Prey species assemblage differences between inner and outer samples of *P. chinensis* nests were investigated using an Analysis of Deviance on a multivariate generalised linear model. A negative binomial distribution was chosen for this analysis based on the dispersion of the residuals. In nests where more than 50 reads of marine origin were detected, the number of reads from commercial seafood and non-commercial sea organisms were analysed in relation to the distance to the sea using the non-parametric Kruskall-Wallis test. All analyses were performed with RStudio and the main packages used for statistical analyses were vegan (Oksanen et al. 2019) and mvabund (Wang et al. 2012).

## RESULTS

### 3.1 Data quality and diet coverage

DNA was successfully amplified and sequenced from all faecal samples. A total of 7,767,737 high quality merged reads were obtained after clean-up of the raw illumina reads. Of these, 7,530,408 could be clustered at 97% identity in 1,436 MOTUs, which were then compared to the NCBI database and analysed thereafter (see supplementary Figure S1). An average of 79,967 DNA reads (± 6,412 s.e.) were obtained per sample. However, the vast majority of those reads (83%) corresponded to fungi, while 13.5% were from terrestrial invertebrate DNA, and 3% of the reads were other taxa, including marine organisms (Figure 2a). Lastly, 0.5% of the reads corresponded to DNA from the wasps themselves. All analyses were performed in R (R Core Team 2016) and the main packages used for statistical analyses were vegan (Oksanen et al. 2019) and mvabund (Wang et al. 2012).

**FIGURE 2.**
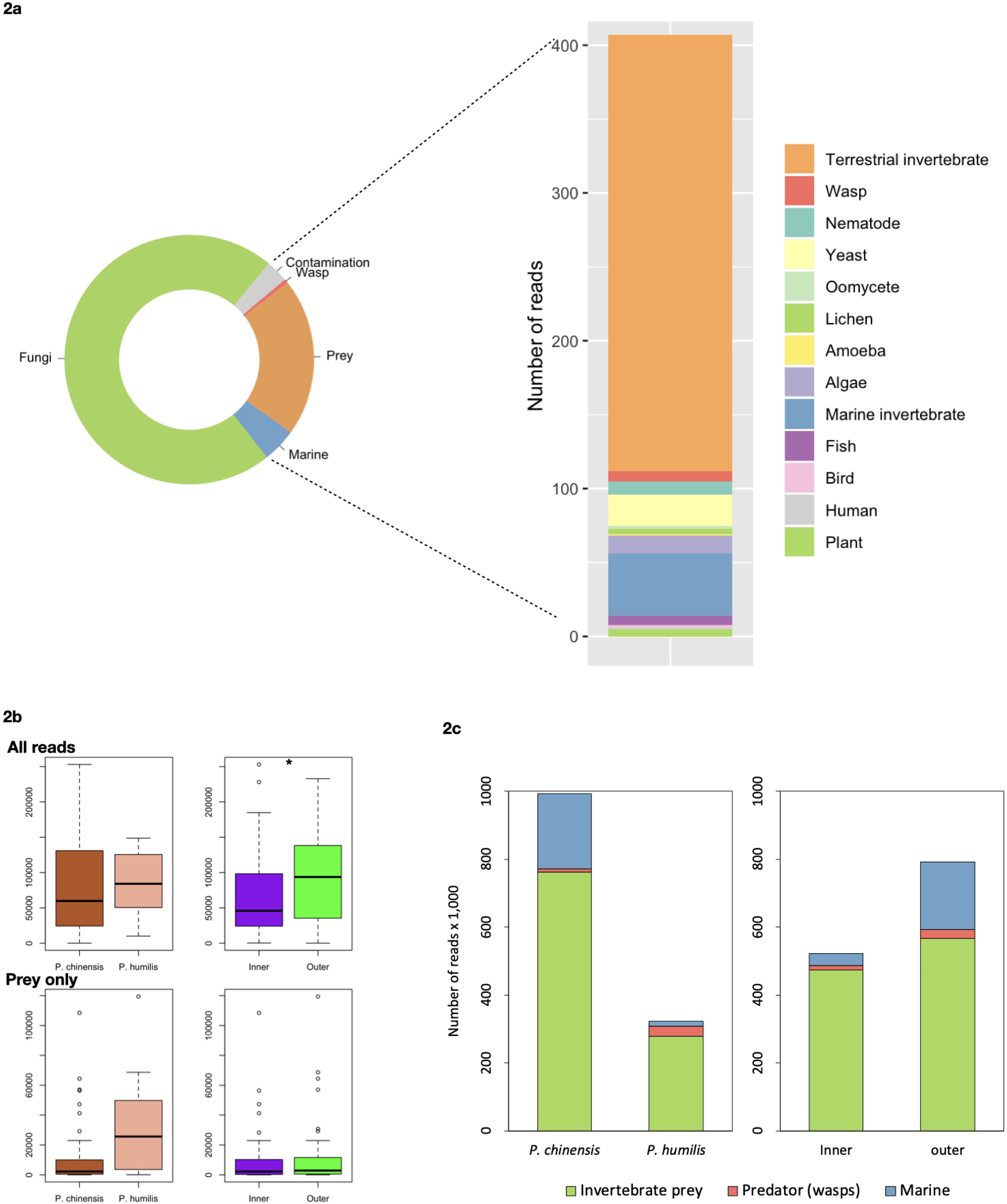
Detection of organisms in the faeces of wasp larvae. 2a. Proportion of MOTUs corresponding to the main categories (left hand-side) and detailed categories excluding fungi (right hand-side) 2b. Number of reads per sample in relation to wasp species (left boxplots), and number of reads in inner versus outer samples (right boxplots) 2c. Total number of reads corresponding to marine, predator and terrestrial invertebrate prey DNA after removal of fungal DNA in both wasp species (left) and in outer versus inner samples (right)

The number of reads per sample was not significantly different between the two species of wasps (KW, *χ* ^2^ = 0.32492, df = 1, p = 0.5687) (Figure 2b). Similarly, there was no significant difference for the number of invertebrate prey reads obtained per sample (*χ* ^2^ = 3.5433, df = 1, p = 0.05978), but a trend could be observed due to an average number of prey reads four times greater in *P. humilis* than in *P. chinensis*. With regard to contamination, no differences in the number of reads per sample were observed between the two wasp species (*χ* ^2^ = 0.029289, df = 1, p = 0.8641) and marine organisms (*χ* ^2^ = 1.3172, df = 1, p = 0.2511).

When comparing inner and outer samples, significantly more reads were obtained from outer samples (Wilcoxon, V = 376, p = 0.02916) (Figure 2b). However, this difference was mainly due to reads considered as contaminants (V = 244.5, p-value = 0.04297) and DNA of marine origin which was five times more abundant in outer samples than in inner samples (V = 299, p = 0.005126) (Figure 2c). In contrast, no differences were found in the number of reads from terrestrial invertebrates when comparing inner and outer samples (V = 524, p = 0.5182). Only 11% of the MOTUs corresponding to terrestrial invertebrates could be identified to the species level, corresponding to 21 species of prey in the diet of *P. chinensis* and eight species in the diet of *P. humilis*. (Table 1, Supplementary figure S2). Therefore, the diet analysis was mainly performed at a higher taxonomic level (order) to provide a greater coverage of the wasps’ diet (i.e. based on all invertebrate MOTUs).

**TABLE 1.**
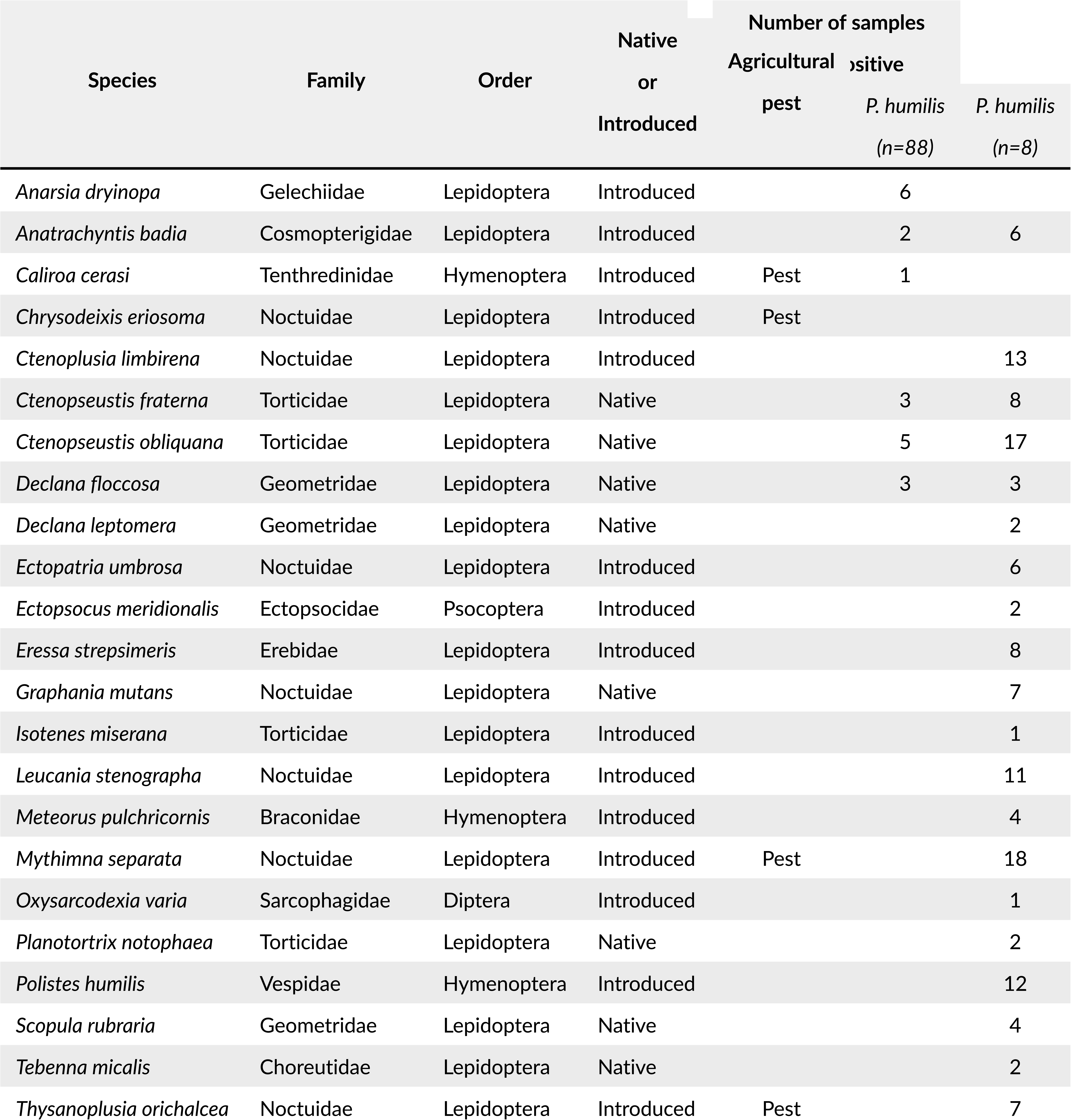
Prey taxa identified at the species level in the diet of *Polistes humilis* and *P. chinensis*

Using 88 samples from *P. chinensis* nests, 260 MOTUs were detected, corresponding to an estimated 91% of the species’ overall terrestrial invertebrate diet (Figure 3). There was no significant difference in the cumulative number of MOTUs detected from inner and outer samples of *P. chinensis* nests (KS, D = 0.25, p = 0.1282) (Figure 3). For *P. humilis*, 141 MOTUs were detected in the eight samples analysed, which only provides a fraction of this species’ diet (Figure 3).

**FIGURE 3.**
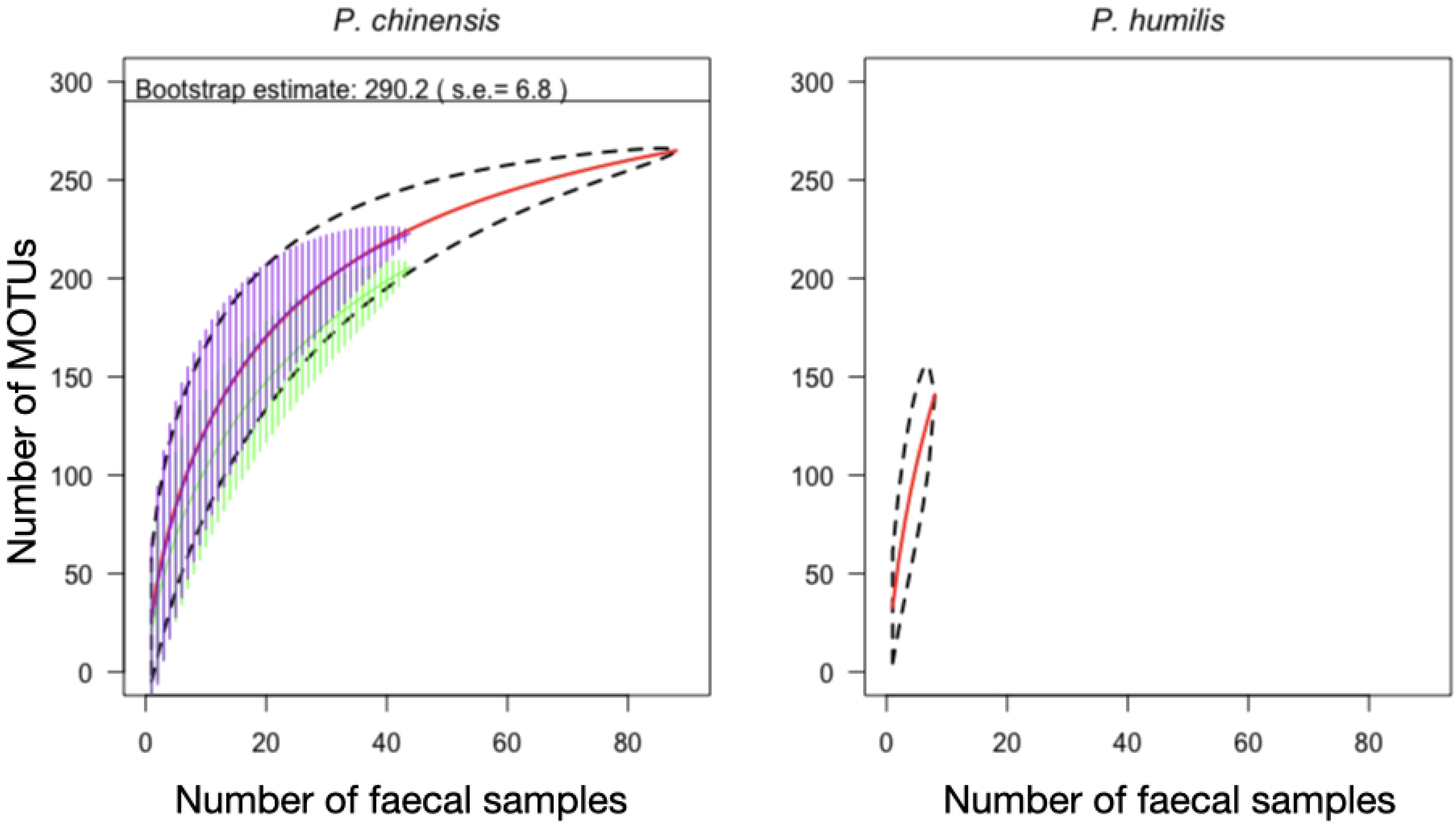
Detection of prey MOTUs in the faeces of wasp larvae. Cumulative curves of number of invertebrate prey MOTUs in relation to number of samples. Left: *P. chinensis*, right: *P. humilis*. Cumulative curves are in red, the area delimited by the dashed lines corresponds to the 95% confidence interval. On the left, the horizontal solid line represents the estimated total number of prey MOTUs in the diet of *P. chinensis*. The purple and green hatched areas correspond to cumulative curves obtained with inner and outer samples respectively.

### 3.2 Prey identification

#### Dev_1,94_

When considering only terrestrial invertebrates, a total of 15 different orders were detected in the diet of *P. chinensis* and nine orders in the diet of *P. humilis*. For both wasp species, the most commonly detected prey belonged to the orders Thysanoptera, Lepidoptera and Acariformes. They were respectively detected in 100%, 74% and 53% of *P. chinensis* samples and in 100%, 88% and 63% of *P. humilis* samples (Figure 4a)

**FIGURE 4.**
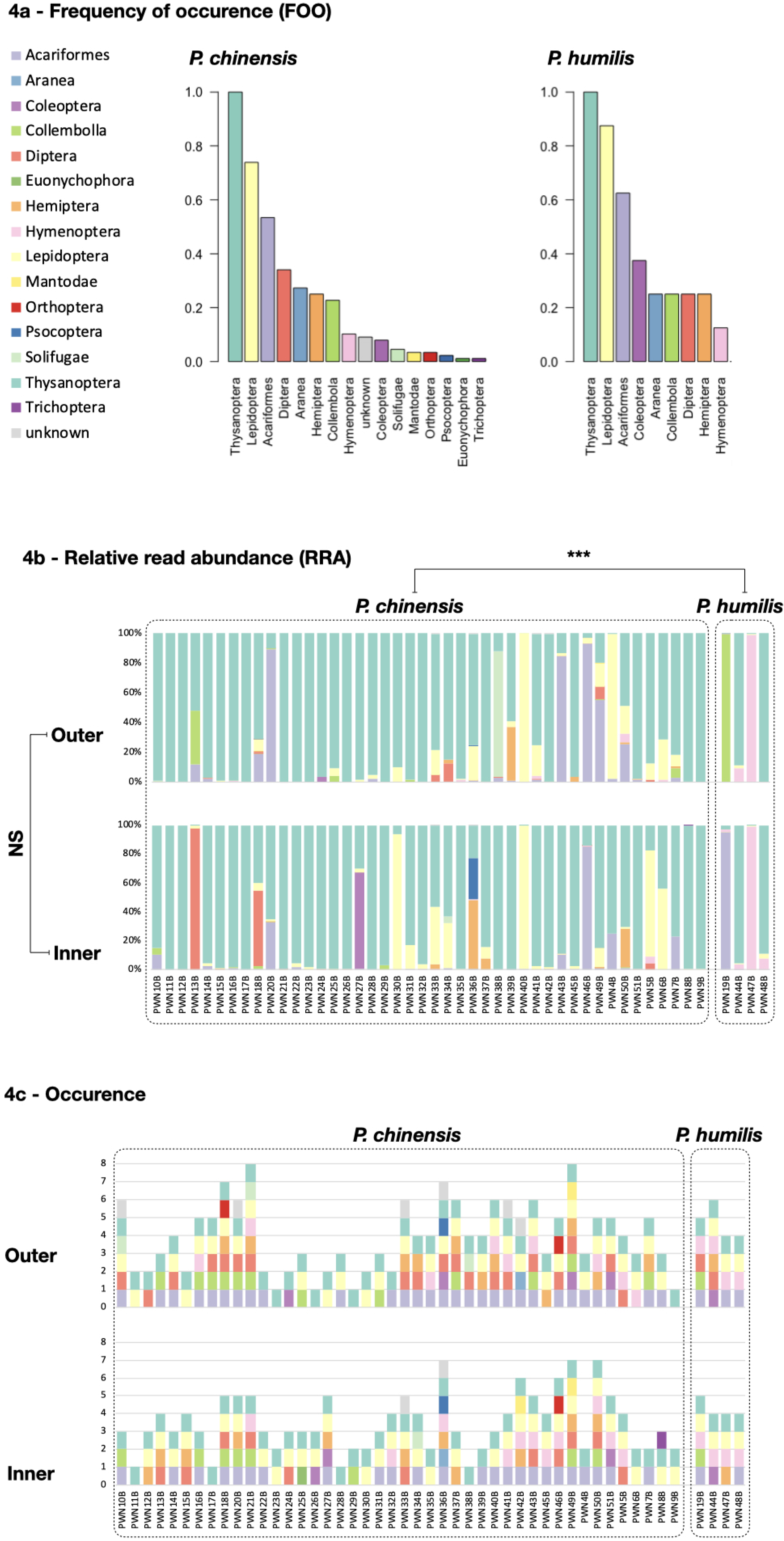
Diet composition of two *Polistes* wasp species from urban and sub-urban areas of Auckland (New Zealand), based on DNA analyses of larval faeces in nests (n=88 for *P. chinensis*; n=8 for *P. humilis*). 4a: Frequency of occurrence of different invertebrate orders in the diet of *P. chinensis* (left) and *P. humilis* (right). 4b: Relative read abundance of prey taxa as measured from each individual faecal sample (i.e. for each individual wasp larvae) in inner and outer cells of the nest. 4c: Occurrence of the different prey genera, in the diet of each individual wasp larvae. See Fig 4a for colour code.

At the MOTU level, the prey community composition (based on RRA) varied with species (Analysis of Deviance: Dev_1,94_ = 713.4, p = 0.002) and to a lesser extent with the location of the samples in the nest (Dev_1,93_ = 576.3 p = 0.032). Similar results were obtained at the order level, with significant differences in prey community composition between wasp species (Dev_1,94_ = 90.18, p = 0.002) (Figure 4b), but only a trend between inner and outer samples (Dev_1,94_ = 51.98, p = 0.083) (Figure 4b). Individual samples contained between 1 and 8 different orders of prey (Figure 4c), and prey richness was significantly higher in outer samples than in inner samples at MOTU level (V = 324.5, p = 0.01845) but only a trend was detected at order level (V = 224, p = 0.05108).

A total of 32 MOTUs could be identified to the species level corresponding to 23 species, most of which were lepidopteran species (18/23) (Table 1). Among these, eight were native species and 15 are considered introduced in New Zealand. The latter includes at least four serious agricultural pests including the cosmopolitan Armyworm (*Mythimna separata* Walker), present in 20% of *P. chinensis* samples.

Over 236,000 reads clustered in 62 MOTUs were identified as marine organisms, mainly corresponding to polychaetes, fish, molluscs and cephalopods (Figure 5). Of these, 93% were recorded in the nests of *P. humilis* (Fig 2c). The vast majority of marine reads were from polychaete worms (Figure 5) and clustered in 31 MOTUs, all matching the genus *Dasybranchus* with 86 to 94% of identity. The only marine MOTU that could be identified to species with confidence (i.e. percentage identity ≥98%) was the squid *Nototodarus gouldi* (McCoy). This species was detected in the nests of both wasps and was present in inner and outer samples. For non seafood organisms, there were no significant differences between nests located close to the sea (< 1 km) and those located further inland (> 1 km), both in terms of number of reads (KW, *χ* ^2^ = 0.88386, df = 1, p = 0.3471) and the number of MOTUs (KW, *χ* ^2^ = 1.5117, df = 1, p = 0.2189). A difference was found however for seafood organisms which produced more reads (KW, *χ* ^2^ = 5.3569, df = 1, p = 0.02064) and more MOTUs (KW, *χ* ^2^ = 9.8955, df = 1, p-value = 0.001657) in nests located more inland (>1 km from the shore) compared to nests closer to shore (Figure 6).

**FIGURE 5.**
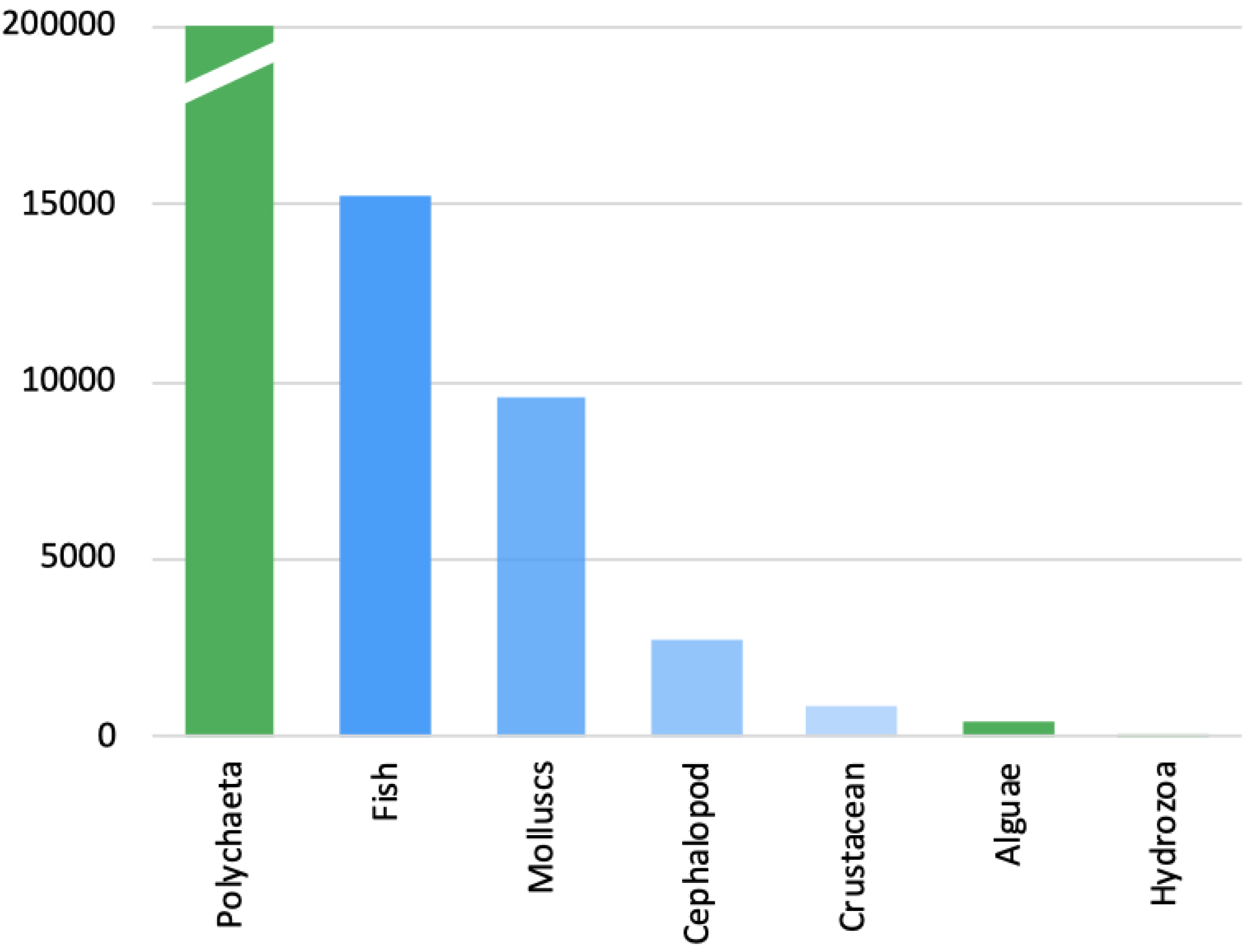
Marine organisms detected in the faeces of paper wasp larvae. Blue bars correspond to potentially commercial sea products, green bars correspond to non-commercial taxa.

**FIGURE 6.**
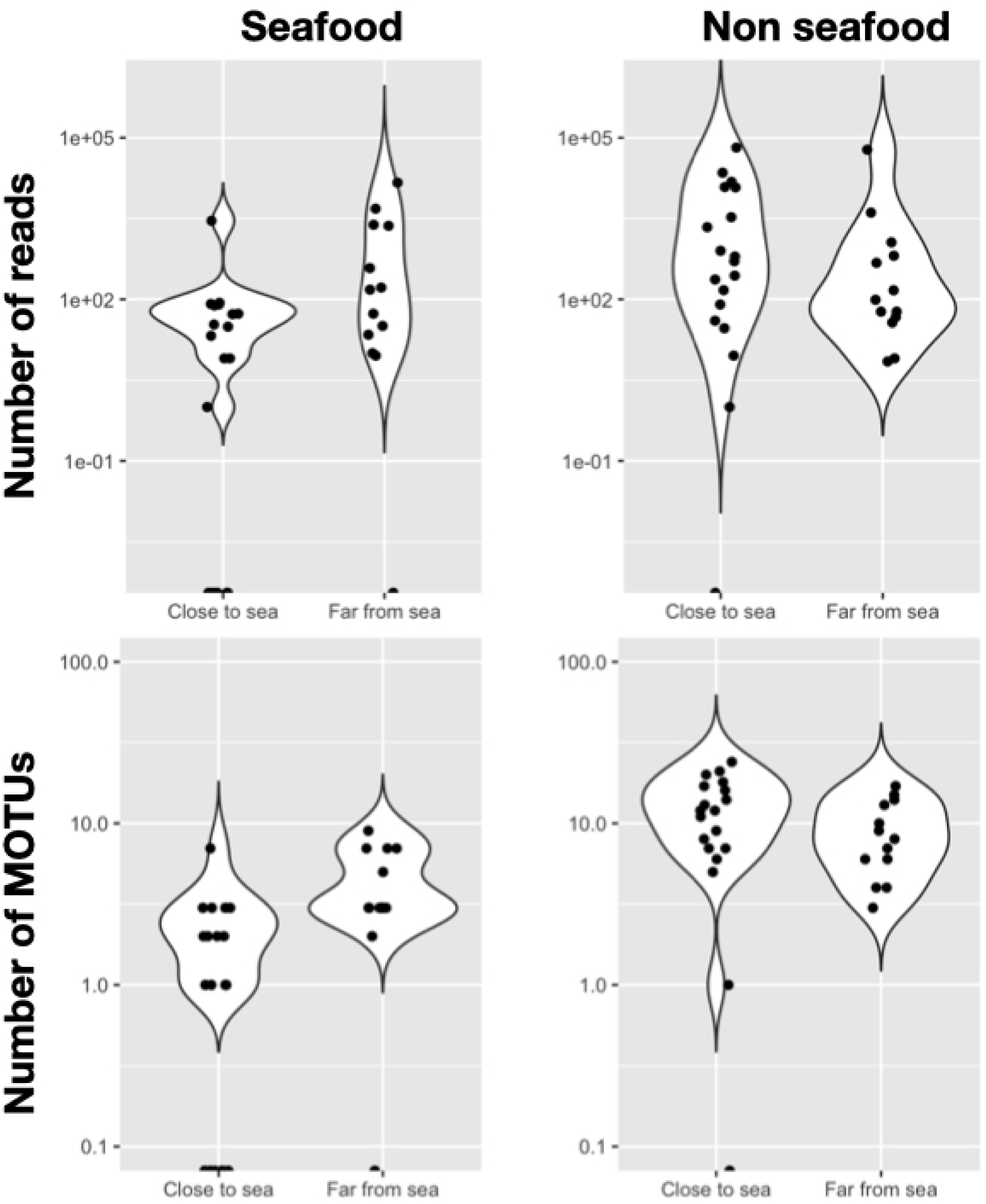
Number of reads for DNA of marine origin in relation to distance to the sea. Samples collected within 1 km of the coast were considered close to sea while samples collected at more than 1 km inland were considered as distant from the sea. Only samples with more than 50 reads from marine origin are represented. Seafood include fish, molluscs, cephalopods, crustaceans and echinoderms, non-seafood include Polychaetes, Tunicates, Hydrozoa and Amoeba.

## DISCUSSION

By analysing prey DNA from faeces accumulated in brood cells, we were able to describe the overall diet of two social Hymenoptera in the urban and peri-urban areas of Auckland, New Zealand. We detected both native species and agricultural pests in the diet of *P. humilis* and *P. chinensis*, and our analysis showed variation between older and more recent faecal samples.

### 4.1 Method efficiency and improvements

Our method led to the successful amplification of prey DNA from paper wasps’ nests and the description of the diet of two polistine wasps in urban and peri-urban areas around Auckland, New Zealand. We used generalist degenerated PCR primers with the aim of amplifying a wide prey spectrum without a priori selection of particular taxa. However, wasp nests appeared to harbour a wide variety of fungi, which were strongly amplified by our primers (1,029 OUT detected). Although we still obtained good numbers of prey reads for most samples, the strong presence of fungal DNA (83% of all sequences generated) suggests a more targeted selection of primers is needed when working on wasp nests. Indeed, the sequencing power lost to fungal DNA would be better used to increase the sequencing depth of prey reads or multiplex more samples in the same sequencing run. Another crucial limitation lies in our inability to identify most prey taxa to species level (Wheeler 2018). The lack of taxonomic resolution encountered here is due to the large proportion of New Zealand invertebrate taxa which are yet to be formally described (Gordon 2010), and the limited availability of sequences for these taxa. Despite the high-resolution power of the chosen primers (Leray et al. 2013), the average percentage identity of prey MOTUs retained in the analyses was only 89.5% (Supplementary Figure S2), which is far from the 98% threshold generally used for identification of invertebrates at the species level.

### 4.2 The case of marine prey

A large number of reads corresponding to species of marine origin were obtained from the faeces of wasps, in particular that of *P. humilis*. To our knowledge, predation of marine organisms by *Polistes* wasps has never been reported before. The great majority of reads belonged to marine worms, probably from the genus *Dasybranchus* Grube, 1850, which includes species present in sand, rocky intertidal regions and shallow waters (e.g. Dean 2016; Mclachlan & Defeo 2018). Such species could be exposed at low tide and may have been targeted by the wasps. However, we found no significant relationship between distance to the sea and number of reads from marine origin. Some of the other marine organisms detected in our samples, such as squid, are unlikely to have been captured by the wasps. While the presence of marine DNA may correspond with actual consumption, it may also reflect insects feeding on carrion washed up by the tide, or feeding on the by-products of human activities (e.g. markets, fishing harbours, food waste). Many of the taxa detected include commercial marine products (fish, crustaceans, cephalopods, echinoderms), which would have been available near harbours or human habitation. Therefore, while these reads may not represent the natural diet of polistine wasps, they do suggest that human activity could strongly influence the diet of some colonies by providing an alternative food source. The greater presence of DNA from marine origin in outer samples (five times more abundant than in inner samples) could suggest a seasonal effect of human fishing activity and/or external contamination by sea spray. Contamination is particularly likely for non-commercial species, mainly represented by polychaete worms, which release large amounts of DNA during swarming events.

### 4.3 Diet composition and ecological role of *Polistes* wasps

The saturation of the MOTU accumulation curve (Figure 3) suggests that the diet of *P. chinensis* was well covered by our analysis. On the other hand, the low number of samples analysed for *P. humilis* did not allow an accurate characterisation of its diet. For the latter species, our results should therefore be taken as indicative only. Our results showed that the two species of wasps displayed a large overlap in their dietary niche (Figure 7) with 147 MOTUs in common. For both species, the diet was dominated, both in terms of relative read abundance and percentage of occurrence, by Thysanoptera (Figure 4a, b), a group known to contain common pests such as thrips. None of the 85 MOTUs corresponding to Thysanoptera (totalling 670,850 reads) could be identified to species level. However, all Thysanoptera MOTUs pointed to the same best hit (with 82 to 92% identity), which was a species in the Phlaeotripidae family. This family is known to comprise a large number of fungus-feeding thrip species that are endemic to New Zealand(Mound & Walker 1986). Another important group, especially in the diet of *P. chinensis*, was Lepidoptera. Identification to the level of species was successful for about a third of the lepidopteran MOTUs (Supplementary Figure S2). The improved identification power for this order is a direct consequence of the significant effort towards describing and barcoding the Lepidoptera fauna of New Zealand (Ball & Armstrong 2006). Lepidopteran prey included three notorious pest species: M. separata, Thysanoplusia orichalcea (Fabricius), and Chrysodeixis eriosoma (Doubleday), found in 41%, 16% and 14% of *P. humilis* samples, respectively. However, the diet of *P. chinensis* also comprised seven native lepidopteran species (mainly Noctuidae, Torticidae and Geometridae). Native and introduced Lepidoptera were also detected in the diet of *P. humilis* and identified to species level. The only pest species that could be identified to species level in the diet of *P. humilis* was the sawfly *Caliroa cerasi* L. It is also interesting to note that the DNA of at least one parasitoid wasp, *Meteorus pulchricornis* (Wesmael), was detected in the nests of *P. humilis*. In New Zealand, this parasitoid is known to attack at least 20 lepidopteran species, both endemic and introduced (Berry & Walker 2004). Although these detections are likely to correspond to secondary predation (given the minute size of the adult parasitoid), by feeding on parasitized caterpillars, *P. humilis* could influence the populations of this parasitoid. These considerations highlight the complexity of the potential impact of *Polistes* wasps predation on New Zealand’s invertebrate fauna.

**FIGURE 7.**
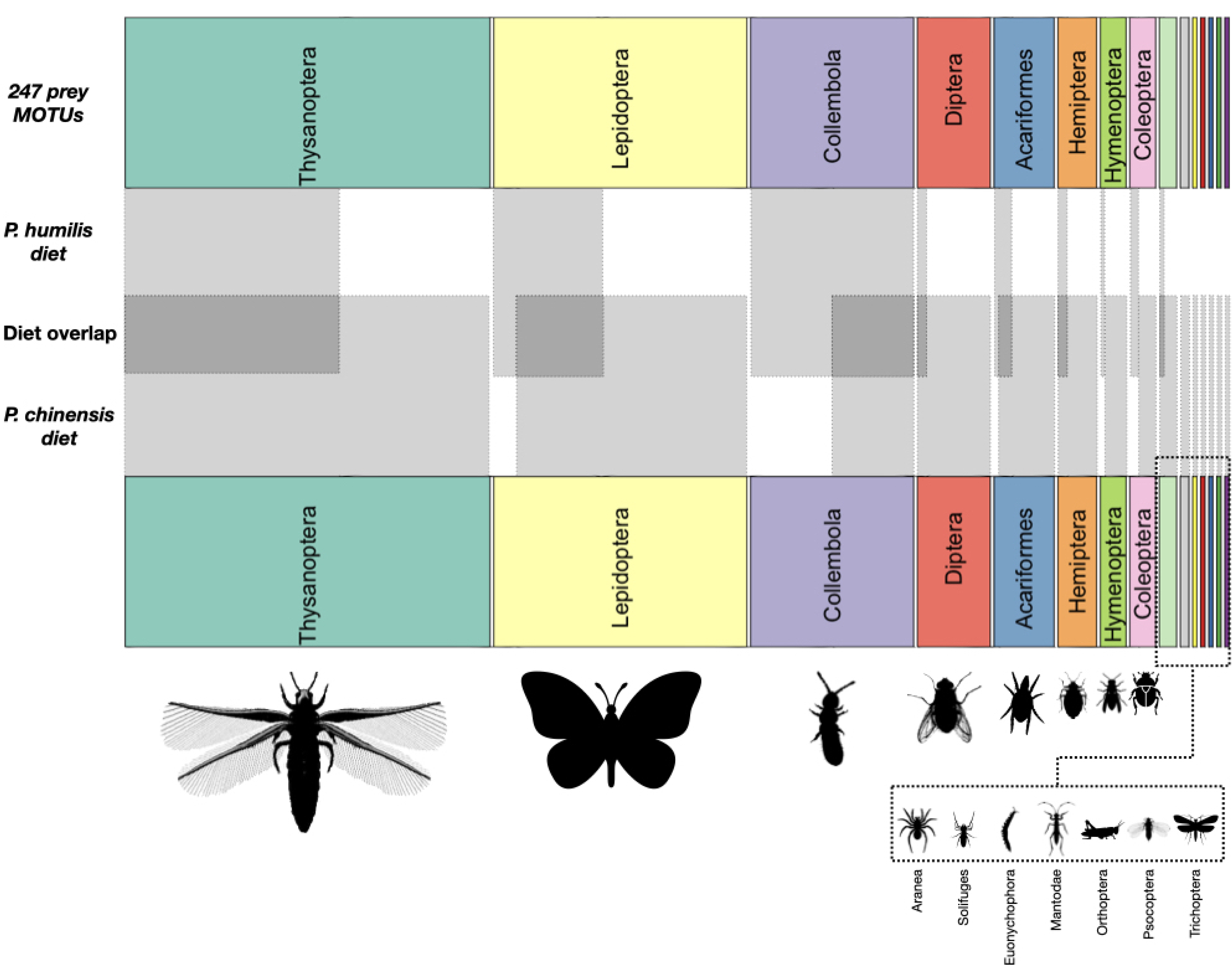
Prey diversity in the diet of *P. humilis* and *P. chinensis* and diet overlap. The horizontal width of the rectangles correspond to the number of MOTU for each order. Light grey areas correspond to MOTUs detected in the diet of each wasp species. Darker grey areas correspond to MOTUs detected in the diet of both wasp species. Most silhouettes were obtained from Phylopic.org.

#### 4.3.1 Seasonal variation in diet

The sizes of collected nests varied over the eleven weeks of collection. Therefore, our samples were not suitable for a well-calibrated seasonal analysis of wasp diet. However, comparing samples taken from inner (older) and outer (younger) layers of the same nest could provide an estimate of how variable the colony diet was within the few weeks necessary to build one or more additional layers of brood cells. The composition of prey communities differed between inner and outer samples, which could reflect variation in prey availability at different times of the year. In addition, we detected greater prey richness in outer (younger) samples, which could indicate that the colony diet diversifies as the number of workers increases. However, this effect could also be explained by the higher degradation of specific prey DNA in inner (older) samples. Therefore, seasonal differences observed must be taken with caution as the rate of DNA degradation was not measured in our study. It is possible that a higher DNA degradation process in older samples conceals part of the early season prey diversity. We also recorded more contamination in samples located on the outer layers that were directly exposed to ambient air, than in samples located on inner layers that were somewhat sheltered inside the nest.

At the very least, the analysis of two faecal samples per nests (inner and outer samples) allowed for a greater coverage of each colony’s diet. However, the number of samples per nest required to accurately estimate the diet of a whole colony may be greater and could vary significantly depending on the size of the nest and the species of interest. It is, for example, known that polistine nests usually contain between 20 to 400 cells, while some *Vespula* species can build significantly larger nests. For example, *Vespula germanica* Fabricius, can build nests comprising over 450,000 cells (Scott 1984) with a colony biomass of up to 600g (Malham et al. 1991). With large colonies in particular, it may be necessary to analyse many cells from the same layer of one nest to test whether a similar diet is provided to all larvae at any given time. Controlled feeding experiments may help to understand how food is divided among the brood. By providing known food items to a captive colony it would also be possible to measure the lifespan of prey DNA inside the nest and precisely measure DNA degradation through time, as well as any potential DNA amplification biases between different types of prey.

Sampling multiple cells from each nests provides more insight into how the colony diet varies over time, and it could identify predatory activity at a much higher resolution through time, potentially mirroring the phenology of the different prey species. However, in *Vespula* wasps, nests are often organised in multiple combs with little if any overlapping cell layers (Rome et al. 2015). In addition, cells remaining on the outer layer can be re-used multiple times by subsequent generations, which means a temporal analysis of the diet using requires precise knowledge of the nest construction.

### 4.4 Concluding remarks

The methodology presented here has the potential to greatly assist in the study of social wasp ecology. Compared to traditional DNA recovery methods, we have developed a method which can provide an overview of the diet of a colony through time based on a single sampling event. If nests are sampled after they are abandoned at the end of the season, it might be possible to uncover a comprehensive record of the colony diet, assuming the degradation of prey DNA remains limited. We hope that this method will be applied to study the ecology of other nest building Hymenoptera, including native and invasive species such as Asian hornets (*Vespa velutina* Lepeletier). Regarding the latter, better knowledge of their diet is essential to measure the ecological impact of their invasion and to ensure the conservation of native invertebrate biodiversity and the ecosystem services they provide (Wycke et al. 2018; Cini et al. 2018).

This study was geographically limited to the urban and peri-urban regions of Auckland. Given this geographic restriction and the relatively low number of nests analysed for *P. humilis*, our results should not be regarded as a comprehensive description of the diet of polistine wasps which are relatively widespread and abundant in many parts of New Zealand (Schmack et al. 2020). Nevertheless, they provide a preliminary insight into the complex role that these species might play in New Zealand. Our results provide field evidence that polistine wasps feed on agricultural pests, supporting the hypothesis that some social wasp species do feed on and may suppress agricultural pests (MacIntyre & Hellstrom 2015a; Southon et al. 2019). Conversely, this study also clearly shows that native New Zealand Lepidoptera are consumed by polistine wasps, thereby illustrating the multifaceted ecological role of these generalist predators.

## 5 AUTHOR CONTRIBUTION

Designed the study: MCL, SB. Collected wasp nests: TS. Performed laboratory analyses: EJD, MCL, SB. Analysed the data, prepared the figures and wrote the first draft of the manuscript: MCL, SB. All authors contributed to the writing of the final manuscript.

## 6 ACKNOWLEDGMENTS

This study was funded by an internal early career research grant obtained my MCL in 2017 at Unitec Institute of Technology (RI17004). Logistical support was provided by the Applied Molecular Solutions Research Focus at Unitec. University of Auckland provided financial support for JRB. We are grateful to the Environmental and Animal Science team at Unitec: Diane Fraser, Mel Galbraith, Dan Blanchon and Lorne Roberts for providing nests and Unitec students for providing first proof of concept during a Conservation Science tutorial (course NSCI 7732, year 2016). University of Auckland provided financial support for JRB.

## 7 SUPPLEMENTARY MATERIAL

**Supplementary Figure S1:**
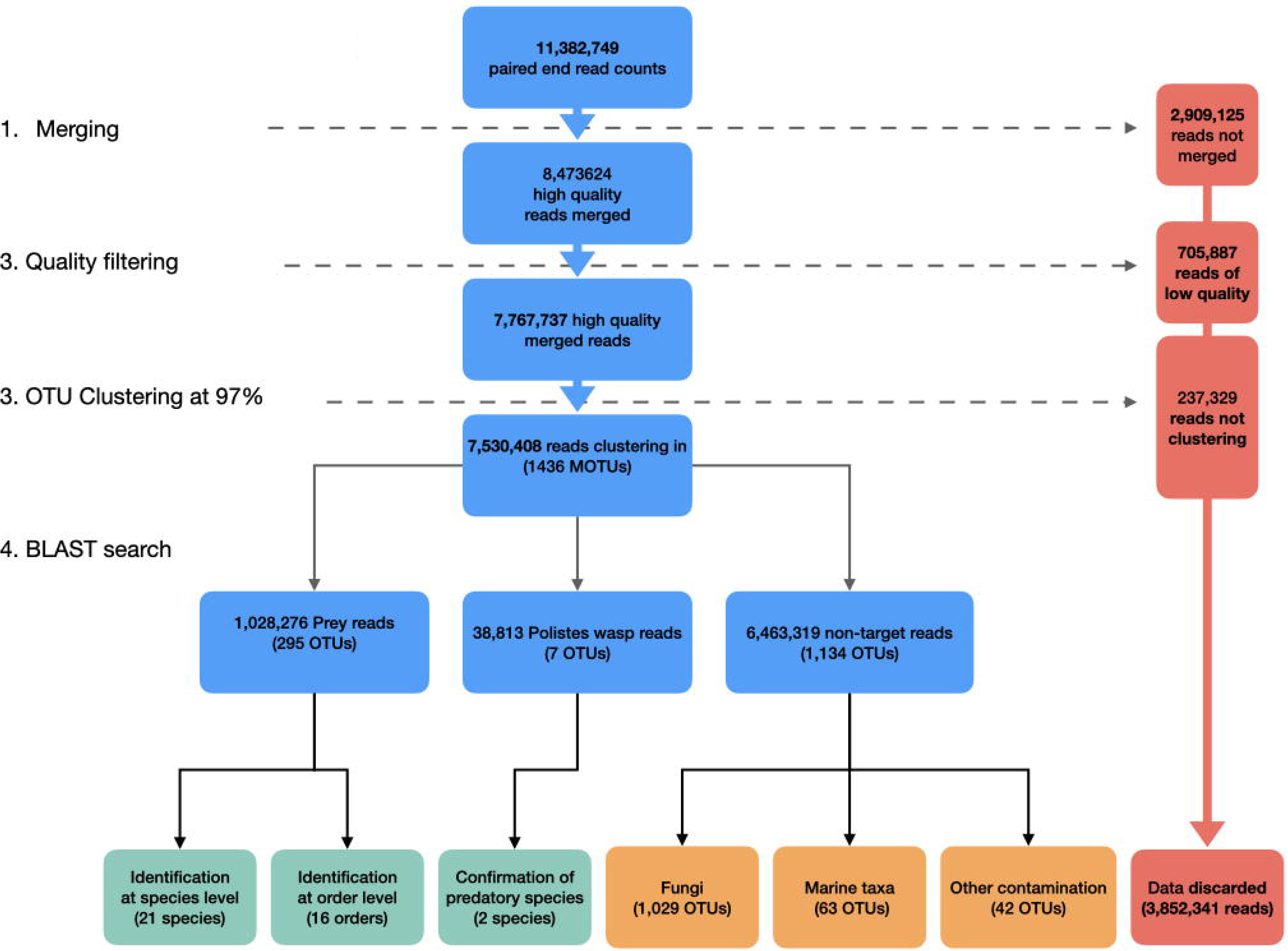
Data processing and number of reads and MOTUs retained or discarded at each step of the bioinformatics analysis

**Supplementary Figure S2:**
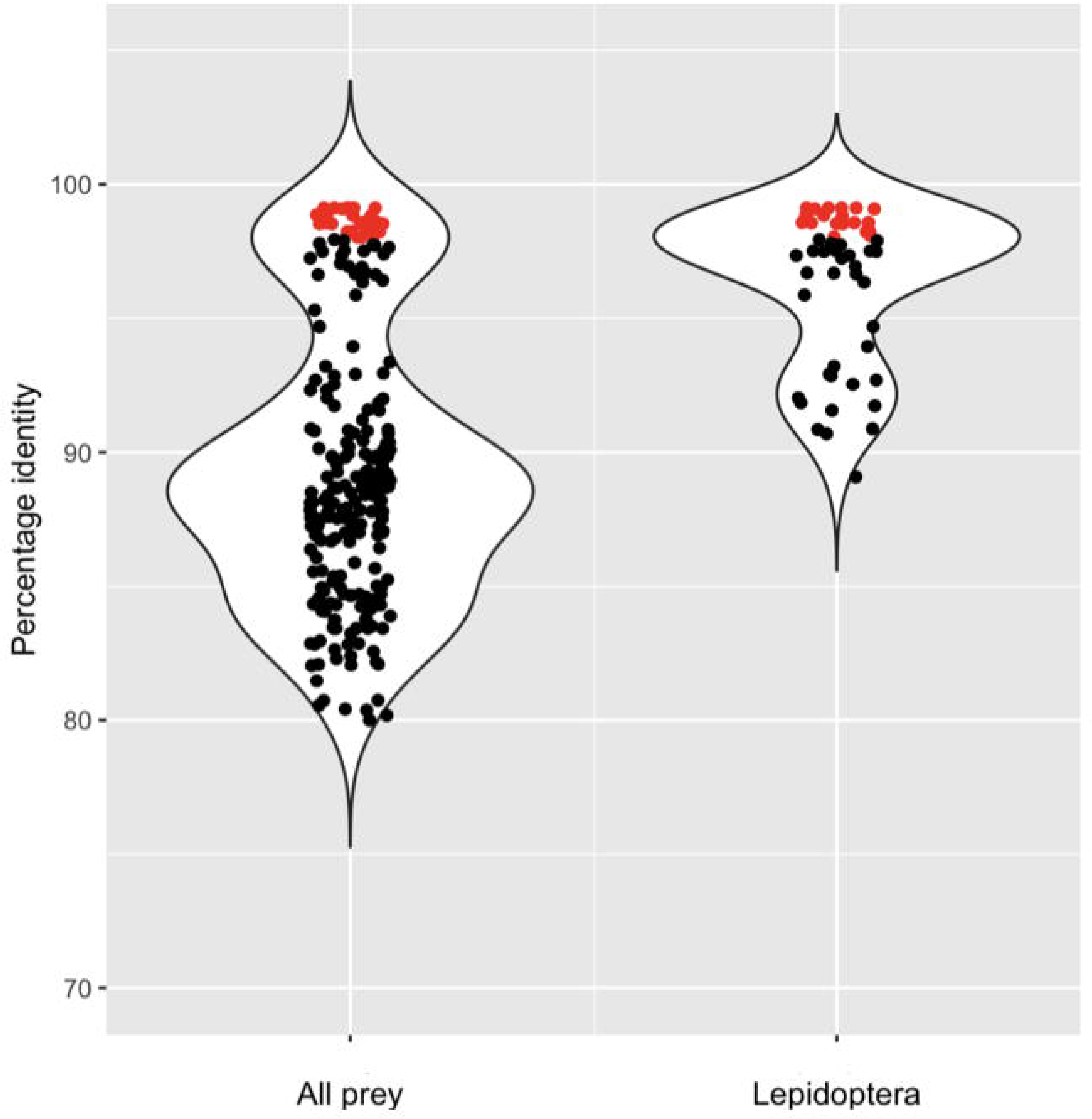
Percentage identity for all prey MOTUs (left) and only Lepidoptera (right). Red dots are MOTUs identified at the species level (i.e. percentage identity ≥98%).

